# JAK inhibitors remove innate immune barriers facilitating viral propagation

**DOI:** 10.1101/2025.02.17.638720

**Authors:** Erlend Ravlo, Aleksandr Ianevski, Marius Nårstad Skipperstøen, Hilde Lysvand, Jørn-Ove Schjølberg, Ole Solheim, Wei Wang, Miroslava Kissova, Marthe Vestvik, Olli Vapalahti, Teemu Smura, Hanna Vauhkonen, Valentyn Oksenych, Friedemann Weber, Mårten Strand, Magnus Evander, Janne Fossum Malmring, Jan Egil Afset, Magnar Bjørås, Denis E. Kainov

## Abstract

Janus kinase (JAK) inhibitors are small-molecule therapeutics that reduce inflammation in autoimmune and inflammatory diseases by modulating the JAK-STAT pathway. While effective in alleviating immune-mediated conditions, JAK inhibitors can impair antiviral defences by suppressing interferon (IFN) responses, potentially increasing susceptibility to viral infections. This study investigates the pro-viral mechanism of JAK inhibitors, focusing on baricitinib, across various cell lines, organoids, and viral strains, including a recombinant Rift Valley fever virus, influenza A virus, SARS-CoV-2, and wild type adenovirus. Our findings demonstrate that baricitinib suppresses transcription of IFN-stimulated genes in non-infected cells (ISGs) which is triggered by type I IFNs produced by infected cells, facilitating viral propagation. The pro-viral effects were influenced by viral load, inhibitor concentration, and structural characteristics of the compound. These results underscore the dual effects of JAK inhibitors: reducing inflammation while potentially exacerbating viral infections. Additionally, the findings highlight opportunities to leverage JAK inhibitors for viral research, vaccine production, and drug screening.

## 1. Introduction

JAK inhibitors are pivotal in managing autoimmune and inflammatory conditions, offering targeted suppression of overactive immune pathways. By modulating the JAK-STAT signalling pathway, these drugs reduce chronic inflammation, improving outcomes in conditions like rheumatoid arthritis, psoriatic arthritis, and atopic dermatitis. Their therapeutic utility extends to hematological disorders and COVID-19 [1]. Currently, ten JAK inhibitors have been approved by the FDA, while numerous investigational compounds are undergoing clinical trials or are in early stages of development [2-4].

JAK inhibitors vary in specificity, with some selectively targeting a single kinase (e.g., upadacitinib for JAK1 and deucravacitinib for TYK2) and others acting more broadly (e.g., baricitinib for JAK1/JAK2 and tofacitinib for JAK1/JAK3). These molecules typically bind to the ATP-binding site of JAKs, and their structural differences influence both their target selectivity and therapeutic applications [5].

Despite their effectiveness in managing inflammation, JAK inhibitors may impair antiviral defence, potentially increasing the risk of viral infections [6]. They are associated with elevated risks of upper respiratory tract infections, viral pneumonia, BK virus viremia and viruria, viral gastroenteritis, and reactivation of viruses such as herpes simplex (HSV), herpes zoster (VZV), and hepatitis B (HBV) [4,7-10]. For example, baricitinib has been linked to an increased incidence of VZV [11]. Patients receiving JAK inhibitors had a significantly higher likelihood of hospitalization or adverse outcomes, including mortality, compared to those treated with interleukin-6 (IL-6) inhibitors or tumor necrosis factor (TNF) blockers [12].

Animal studies provided compelling evidence of the impact of JAK inhibition on viral infections. For example, JAK inhibition during the early phase of SARS-CoV-2 infection has been shown to exacerbate kidney injury in mice by suppressing antiviral responses [13]. Similarly, tofacitinib treatment worsened outcomes in HSV infection models [14]. In vitro studies have also shown that JAK inhibitors could suppress antiviral responses and enhance replication of VSV, RSV, IAV, MeV, MuV, ZIKV, HHV-6A, ReoV [15-21]. Collectively, these findings suggest that JAK inhibitors may increase susceptibility to infections, though the detailed mechanism behind this adverse effect is not fully understood.

It is important to investigate the effects of JAK inhibitors on other viral infections, such as Rift Valley fever virus (RVFV). RVFV is a zoonotic pathogen that can infect humans, causing a wide range of clinical outcomes, from mild febrile illness to severe complications including hepatitis, hemorrhagic fever, and meningoencephalitis [22]. The primary organs affected by RVFV include the liver, lungs, eye and brain. Among these, neurological involvement, particularly encephalitis and other forms of brain damage, is considered a major contributor to mortality [23]. Understanding how JAK inhibitors influence the course of RVFV infection, especially in the context of neuroinflammation and viral replication in the central nervous system, may provide novel insights into pathogenesis of the disease.

Here we bridge this gap by systematically elucidating the pro-viral effect of JAK inhibitors, identifying key factors such as drug concentration, chemical structure, and viral load that influence their impact on antiviral immunity. We showed how immunosuppressive properties of Jak inhibitors can compromise antiviral defenses, presenting risks during active infections.

## 2. Materials and Methods

### 2.1. Small molecules, viruses, cell and organoids

Abrocitinib (HY-107429), AT9283 (HY-50514), baricitinib (HY-15315), delgocitinib (HY-109053), fedratinib (HY-10409), filgotinib (HY-18300), itacitinib (HY-16997), oclacitinib (HY-13577), pacricitinib (HY-16379), ruxolitinib (HY-50856), tofacitinib (HY-40354), and upadacitinib (HY-19569) were obtained from MedChemExpress. To obtain 10 mM stock solutions, compounds were dissolved in dimethyl sulfoxide (DMSO; Sigma-Aldrich, Germany) or water. The solutions were stored at -20 °C until use.

The recombinant rRVFVΔNSs::Katushka expressing the far-red fluorescent protein Katushka instead of the NSs protein (rRVFV) was generated as described previously [24]. The recombinant influenza A/PR/8-NS116-GFP strain (rIAV), expressing GFP instead of the effector domain of NS1, was generated as described previously [25]. Recombinant SARS-CoV-2/Wuhan-mCherry (rSARS-CoV-2), expressing mCherry, was generated as described previously [26]. Adenovirus (AdV) was derived from a nasopharyngeal swab sample collected at St. Olavs Hospital.

Human adenocarcinomic alveolar basal epithelial cells (A549, CCL-185), hTERT-immortalized retinal pigment epithelial cells (RPE, CRL-4000), human lung adenocarcinoma epithelial cells (Calu-3, HTB-55), Madin-Darby canine kidney cells (MDCK, CCL-34), monkey Vero-E6 cells (Vero C1008, CRL-1586) cell lines were obtained from the ATCC. ACE2-expressing A549 cells (A549-ACE2) were from [27]. A549, and Vero-E6 cells were grown in Dulbecco’s Modified Eagle’s medium (DMEM; Gibco, Paisley, Scotland) supplemented with 100 U/mL penicillin and 100 μg/ml streptomycin mixture (pen/strep; Lonza, Cologne, Germany), 4.5 g/L (=25 mmol/L) glucose, 1 mM L-glutamine, and 10% heat-inactivated fetal bovine serum (FBS; Lonza, Cologne, Germany). RPE and Calu-3 cells were grown in DMEM-F12 supplemented with 10% FBS, pen/strep. All cells were cultured at 37ºC with 5% CO2, 95% humidity and passaged using 0.05% (v/v) Trypsin/EDTA (Gibco). Cells were tested mycoplasma negative throughout the work (MycoAlert Mycoplasma Detection Kit, Lonza).

rRVFV and rSARS-CoV-2 were amplified in a monolayer of Vero-E6 cells in growth media containing 1% FBS. rIAV was amplified in a monolayer of MDCK cells in virus growth media containing 0.2% bovine serum albumin (BSA; Sigma) and 0.2 μg/mL TPSK-Trypsin (Sigma). Virus stocks were stored at −80 °C.

Induced pluripotent stem cells (iPSCs) were used to generate retinal organoids as described previously [28] [29]. Glioblastoma organoids have been produced following the protocol by Jacob *et al* [30].

### 2.2. Drug treatment and virus infection of cells and organoids

Approximately 4 × 10^4^ cells were seeded in each well of 96 well/plate. After 24 h small molecules were added. 15 min later cells were infected. The infection of A549 and Vero-E6 cells with rRVFV was done in DMEM-based medium containing 1% FBS. The infection of RPE and Calu-3 cells with rRVFV was done in DMEM-F12-based media containing 1% FBS. The infection of A549 cells with AdV was done in DMEM-based medium containing 1% FBS. The infection of A549 cells with rIAV was done in DMEM-based media containing 0.2% BSA and 0.2 μg/mL TPSK-Trypsin. The infection of A549-ACE2 cells with rSARS-CoV-2 was done in DMEM-based medium containing 1% FBS.

40-days old retinal organoids were transferred from petri dishes to an ultralow attachment 96-well plates. One organoid was placed in each well in 96-well plate, with a total of 100 µl of media per well. 5 µM baricitinib or vehicle were added. Virus at multiplicity of infection (moi) 0.1 or mock was added after 15 min.

### 2.3. Cell and organoid imaging, viability and death assays

The cell and organoids were imaged using Evos FL Auto imaging system (Invitrogen). Cell viability and cytotoxicity were assessed using the CellTiter-Glo (CTG) and CellTox Green (CTxG) assays (Promega, Cat. Nos. G9241 and G8741, respectively). Measurements were performed on a FluoStar Omega plate reader (BMG Labtech), followed by quantification of luminescence and fluorescence signals.

### 2.4. Calculation of drug sensitivity scores

Drugs were added to cells in 3-fold serial dilutions. Cells were infected with viruses (moi 0.1) or mock. The cell viability was measured using CTG assay after 48 h of infection. A drug sensitivity score (DSS) was calculated as a normalized version of the standard area under dose–response curve (AUC), with the baseline noise subtracted, and the normalized maximal response at the highest concentration (often corresponding to off-target toxicity):

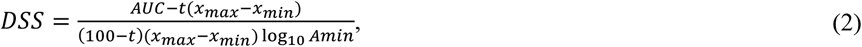

where activity threshold *t* equals 10%, and DSS is in the 0-50 range [31-34]. The difference (ΔDSS) between DSS (virus) and DSS (mock) was also calculated.

### 2.5. Protein profiling

The proteome profiler human XL cytokine array kit (ARY022B), proteome profiler human apoptosis array kit (ARY009), and proteome profiler phospho-kinase array kit (ARY003C) from R&D Technologies were used to profile extracellular cytokines, intracellular proteins involved in apoptosis, and the phosphorylation status of proteins associated with various signaling pathways. All arrays were performed according to the manufacturer’s protocols.

### 2.6. RNA sequencing

Total RNA was isolated from samples using NAxtra kit following the manufacturer’s protocol [35]. For library preparation, mRNA was enriched from the total RNA using oligo-dT beads to select for polyadenylated transcripts. The enriched mRNA was fragmented into smaller pieces, which were used for cDNA synthesis. First-strand cDNA synthesis was performed using random primers, followed by second-strand synthesis incorporating dUTP in place of dTTP. The resulting cDNA fragments were end-repaired and adenylated at the 3′ ends, after which adaptors were ligated. Libraries were then amplified via polymerase chain reaction (PCR) and assessed for quality. The amplified libraries were circularized and further amplified to produce DNA nanoballs (DNBs). Sequencing was performed on the DNBSEQ Technology platform with a paired-end read length of 150 bp (PE150), using the DNBSEQ Eukaryotic Strand-specific mRNA library, as specified for DNBSEQ Eukaryotic Strand-specific Transcriptome Resequencing. Raw sequencing data were processed to remove low-quality sequences and adapter contamination. Reads containing 25% or more adapter sequence (allowing up to 2 mismatches) were discarded. Reads shorter than 150 bp were removed, as were those with an N content of 0.1% or more. Reads with stretches of a single nucleotide exceeding 50 bp were excluded. Additionally, reads where bases with a quality score below 20 constituted 40% or more of the sequence were filtered out. The resulting high-quality reads were retained for downstream analysis, with quality scores reported using the Phred33 system.

RNA-seq reads were aligned using STAR (Spliced Transcripts Alignment to a Reference) version 2.7.11b to the reference human GRCh38 genome. Gene-level read counts were obtained using STAR’s built-in-quantMode GeneCounts function. Gene symbols and descriptions were assigned using the biomaRt package (v 2.58).

Differential gene expression was performed using DESeq2 (v1.34.0). Genes with an adjusted p-value < 0.05 and absolute log2 fold change > 1.5 were considered significantly differentially expressed. Pathway enrichment analysis was conducted using the clusterProfiler R package (v. 4.8.3) to identify overrepresented biological pathways among differentially expressed genes. Pathways with adjusted p-value < 0.05 were considered significantly enriched. [36]. Results were visualized as bubble plots, where the x-axis represented enriched pathways, the y-axis showed experimental comparisons, bubble size indicated the gene count (number of genes in each pathway), and bubble colour intensity reflected statistical significance (−log10(adjusted p-value)).

The RNA-seq data have been deposited in the NCBI Gene Expression Omnibus (GEO) under accession number GSE292720.

### 2.7. qPCRs

Quantitative PCR (qPCR) was performed on the CFX Connect Real-Time PCR Detection System (Bio-Rad) using qScript™ 1-Step Virus ToughMix (Quantabio) and virus-specific primers and TaqMan probe (Table S1). Quantitative RT-PCR (RT-qPCR) was performed using host-specific primers (Table S2).

## 3. Results

### 3.1. Potential pro-viral mechanism of action of the JAK inhibitors

We hypothesized that certain JAK inhibitors, including approved ones (Fig. S1), may suppress the IFN response and facilitate the progression of viral infections through a straightforward mechanism. In this mechanism, viral infections stimulate the synthesis of IFNs in host cells, which are subsequently released into the extracellular space. IFNs then bind to receptors on the surface of uninfected cells and activate the JAK/STAT signaling pathway, leading to the synthesis of ISGs that protect the cells from potential infection. However, JAK inhibitors disrupt this protective response in uninfected cells, allowing viruses to infect new cells unimpeded and spread throughout the tissue (Fig. 1).

**Figure 1.**
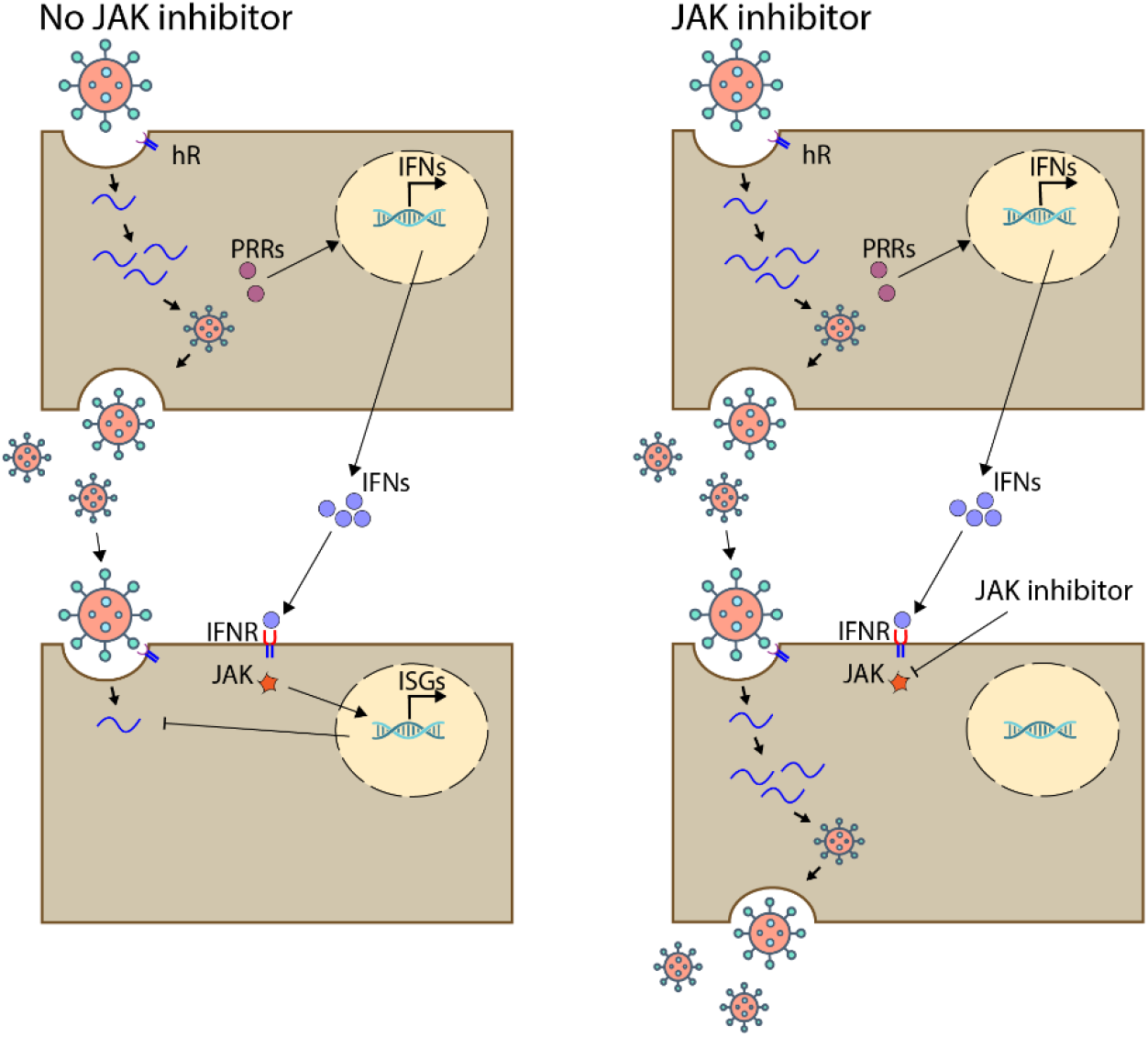
Potential pro-viral mechanism of action of the JAK inhibitors. Schematics showing how viral infections induce type I interferon (IFN) synthesis in infected cell and its secretion. Secreted IFN triggers JAK/STAT pathway that activates ISG production, creating an antiviral state in uninfected cells (left panel). JAK inhibitors disrupt JAK/STAT pathway, removing innate immune barriers and enabling the amplification of viral particles in previously uninfected cells (right panel).

### 3.2. Baricitinib enhances rRVFV infection in A549 cells

To prove the hypothesis, we first assessed the impact of baricitinib on rRVFV virus infection. A549 cells were treated with 5 µM baricitinib or a vehicle control (0.01% DMSO) and infected with the virus at a moi of 0.1. Fluorescent microscopy showed a time-dependent increase in the expression of the fluorescent Katushka protein in baricitinib-treated cells compared to the vehicle control (Fig. 2a, b). Notably, the treated cells remained viable after 36 hpi (Fig. 2c).

**Figure 2.**
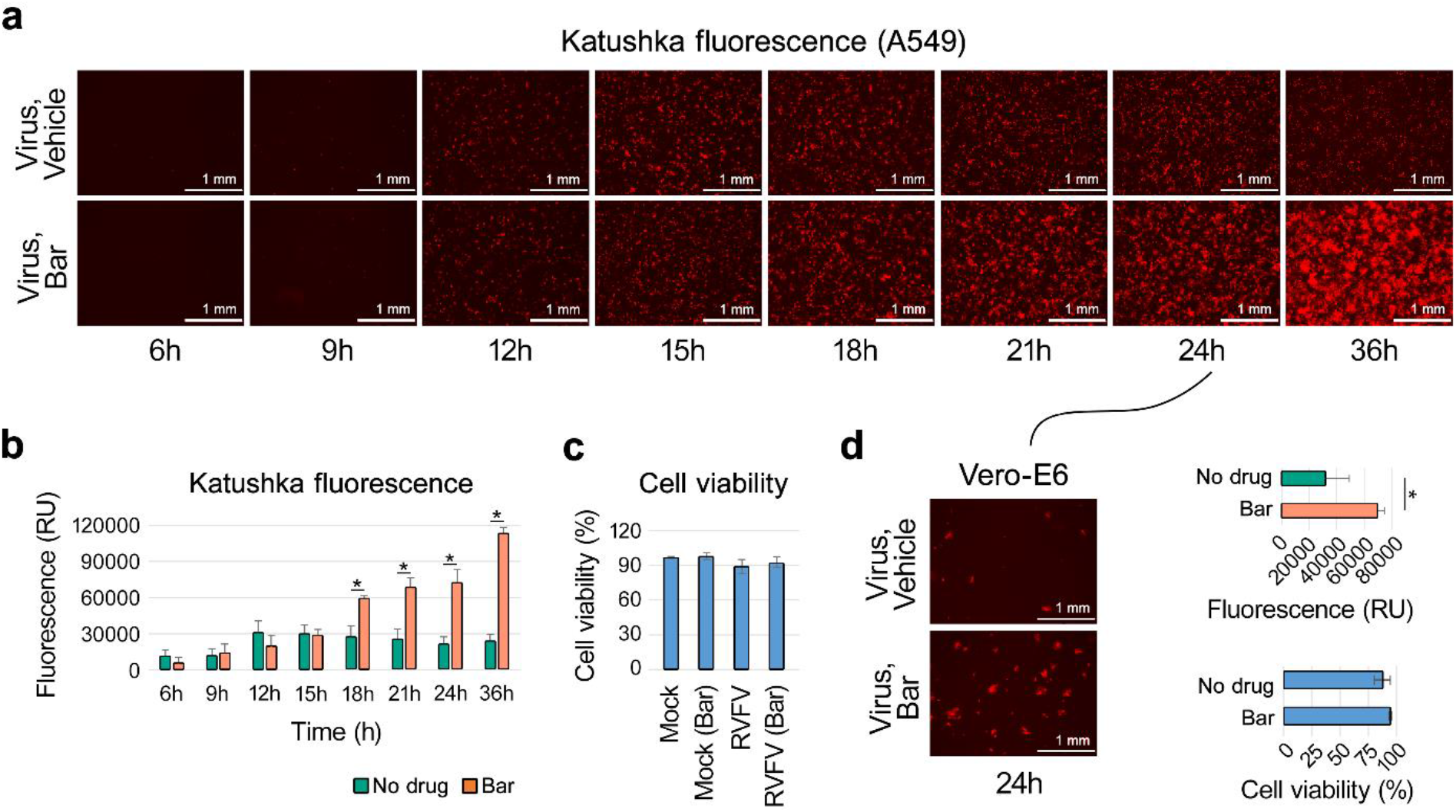
Baricitinib enhances the progression of rRVFV infection in A549 cells. (**a, b**) A549 cells were treated with either a vehicle or 5 µM baricitinib and infected with the rRVFV virus (moi 0.1). Images of the infected cells were captured, and fluorescence of far-red signal intensity was quantified. The fluorescence signal was normalized and expressed as Mean ± SD (n = 3). (**c**) After 36 hours, cell viability of virus- and mock-infected, vehicle- and baricitinib-treated A549 cells was assessed using a CTG assay (mean ± SD, n = 3). (**d**) Supernatants from vehicle- and baricitinib-treated A549 cells were collected after 24 h of virus infection, diluted 1:10000 and applied to Vero-E6 cells. Katushka reporter protein expression and cell viability were evaluated as described in (a–c). Statistical significance is indicated as * p < 0.05, determined by Wilcoxon rank-sum test with alternative hypothesis ‘greater than’.

We next evaluated the production of infectious virus particles. Supernatants from vehicle- and baricitinib-treated A549 cells were collected 24 hours post-infection, diluted 1:1000, and applied to IFN-I-deficient Vero-E6 cells [37,38], which are not sensitive to baricitinib during virus propagation (Fig. S2a). Analysis of reporter protein expression and cell viability demonstrated that baricitinib treatment resulted in the production of higher levels of infectious virus particles (Fig. 2d).

We performed a time-of-addition experiment in A549 cells using the virus (Fig. S3). The results showed that the baricitinib can be added between 0 and 6 hours (corresponds to time of IFN-induction) after infection to promote viral replication. These findings indicate that baricitinib enhances virus replication supporting our initial hypothesis.

### 3.3. The pro-viral effect of JAK inhibitors is influenced by initial rRVFV load, concentration and chemical structure of the small molecules

To investigate the impact of viral load on the pro-viral effects of baricitinib, A549 cells were treated with either vehicle or baricitinib and subsequently infected with rRVFV at varying moi. After 24 h, fluorescent and bright-field microscopy images were captured, and Katushka reporter protein expression along with cell viability were measured. The results demonstrated that baricitinib exhibited a substantial pro-viral effect at moi below 0.1 (Fig. 3a,c). These findings support the hypothesis that at least one round of viral replication is necessary to induce type I IFN production in infected cells, which allows baricitinib to block the IFN-mediated antiviral state in uninfected cells, thereby promoting subsequent rounds of viral replication.

**Figure 3.**
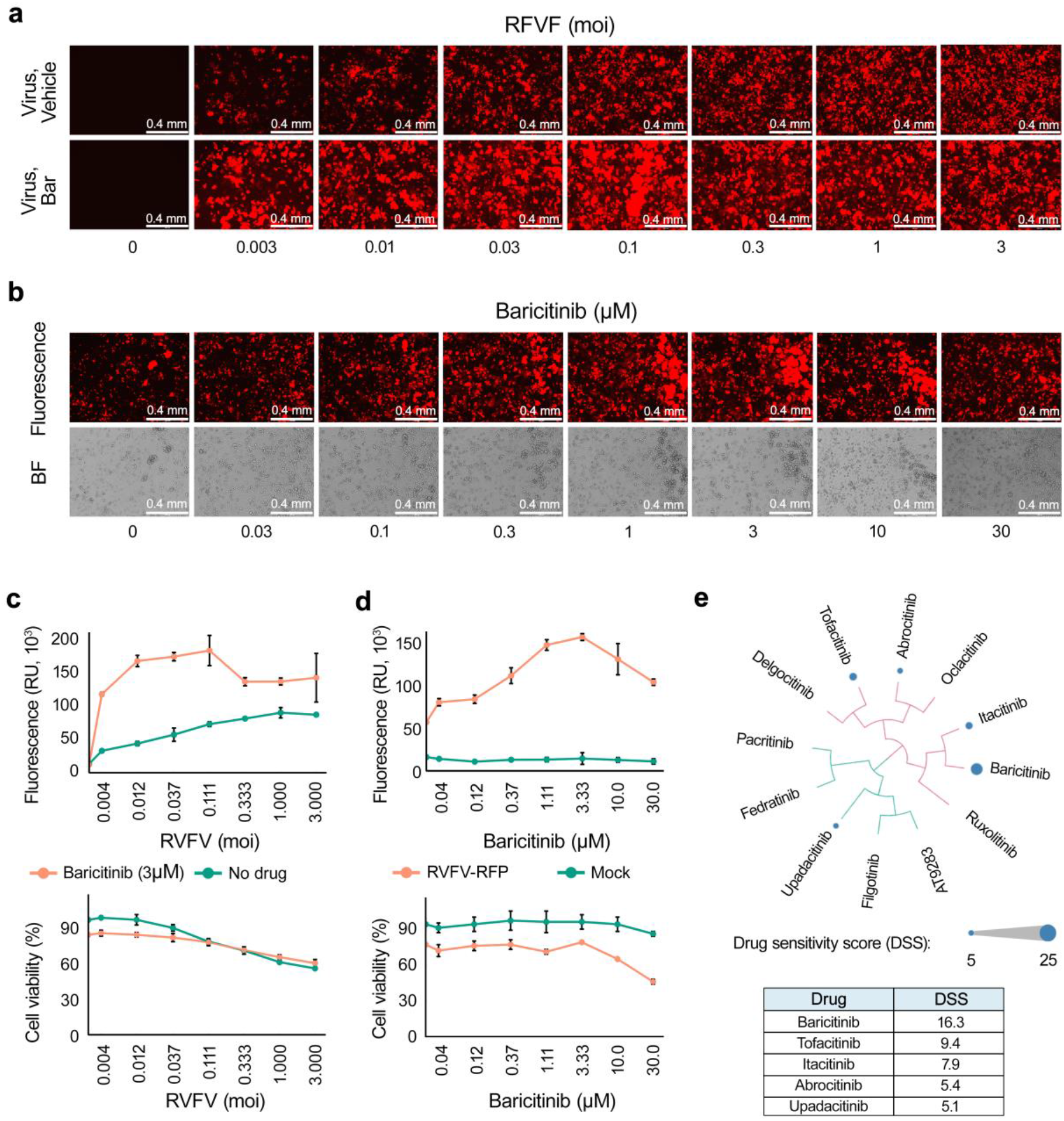
The pro-viral effect of JAK inhibitors is influenced by initial virus load, concentration and chemical structure of small molecules. (**a, c**) A549 cells were treated with either a vehicle or 5 µM baricitinib and infected with the rRVFV at varying moi. After 24 hours, fluorescent and bright-field images were captured, Katushka reporter protein expression and cell viability were quantified (mean ± SD, n = 3). (**b, d**) A549 cells were treated with vehicle or different concentrations of baricitinib and infected at an moi of 0.1. After 24 hours, fluorescence and bright-field images were captured, and Katushka reporter protein expression and cell viability were analyzed (mean ± SD, n = 3). (**e**) A549 cells were treated with varying concentrations of different JAK inhibitors. DSS were calculated and visualized as bubbles in a SAR diagram. Bubble sizes represent DSS values, and compound similarities were determined using ECFP4 fingerprints and visualized through the D3 JavaScript library. The table (bottom panel) lists the exact DSS values for the five most active JAK inhibitors, where DSS (Drug Sensitivity Score) is a normalized area under the dose-response curve (AUC, see methods)..

To evaluate the concentration-dependent effects, A549 cells were treated with vehicle or increasing concentrations of baricitinib and infected with the virus at an moi of 0.1. After 24 h, fluorescence and bright-field images were captured, and both Katushka reporter protein expression and cell viability were analyzed. Baricitinib displayed a dose-dependent enhancement of viral replication up to 10 μM concentration, when the drug became toxic (Fig. 3b,d).

The effects of 11 additional JAK inhibitors were subsequently assessed in A549 cells under conditions similar to those used for baricitinib. DSS were calculated for each compound and visualized as bubbles in a structure-activity relationship (SAR) diagram (Fig. 3e), where bubble sizes correspond to DSS values. Among the tested inhibitors, upadacitinib, tofacitinib, itacitinib, and abrocitinib, in addition to baricitinib, demonstrated the ability to promote viral replication, whereas seven other JAK inhibitors did not exhibit this pro-viral effect at the tested concentrations (Fig. S4).

Structural analysis revealed that the five active compounds shared a heterocyclic aromatic core, which serves as the molecular framework for specific interactions with the ATP-binding site of the JAK kinase domain (Fig. S5). These findings suggest that the pro-viral effects of JAK inhibitors are influenced by both their chemical structure and concentration, with the heterocyclic aromatic core playing a pivotal role.

We screened 300 compounds from our in-house broad-spectrum antiviral (BSA) library in combination with baricitinib [39,40]. Our pilot screen revealed that A-1155463 and ABT-737 (both pro-apoptotic), as well as ozanimod (a sphingosine-1-phosphate receptor modulator), pimodivir (a PB2 inhibitor targeting influenza virus replication), and tiplaxtinin (a selective inhibitor of plasminogen activator inhibitor-1) enhanced the expression of the Katushka protein in rRVFV-infected A549 cells (Fig. S6) [41-44]. By contrast, anisomycin (a protein synthesis inhibitor) suppressed eRVFV-mediated Katushka expression. These findings suggest that other compounds might further amplify the pro-viral effects of Jak inhibitors such as baricitinib.

### 3.4. Baricitinib attenuates transcription of IFN-stimulated genes and modulates apoptotic, and signaling pathways in A549 cells during rRVFV infection

To investigate the effect of baricitinib on the host immune response to RVFV infection, A549 cells were treated with 5 µM baricitinib or vehicle, and infected with rRVFV or mock. RNA sequencing analysis revealed significant transcriptional changes during RVFV infection and baricitinib treatment. Differential expression analysis presented as a heatmap in Figure 4a shows that rRVFV infection strongly upregulated key interferon-stimulated genes (ISGs) such as MX1, MX2, IFIT3, and IFI44L, compared to mock infection. Baricitinib treatment showed minimal effects on gene expression in uninfected cells but significantly attenuated the ISG response in RVFV-infected cells. The detailed statistical analysis of these differentially expressed genes is presented in volcano plots in Figure S7, which further illustrates the magnitude and significance of these transcriptional changes. Pathway enrichment analysis confirmed that rRVFV activated antiviral and immune signaling, while baricitinib suppressed activation of type I IFN pathway (Fig. 4b).

**Figure 4.**
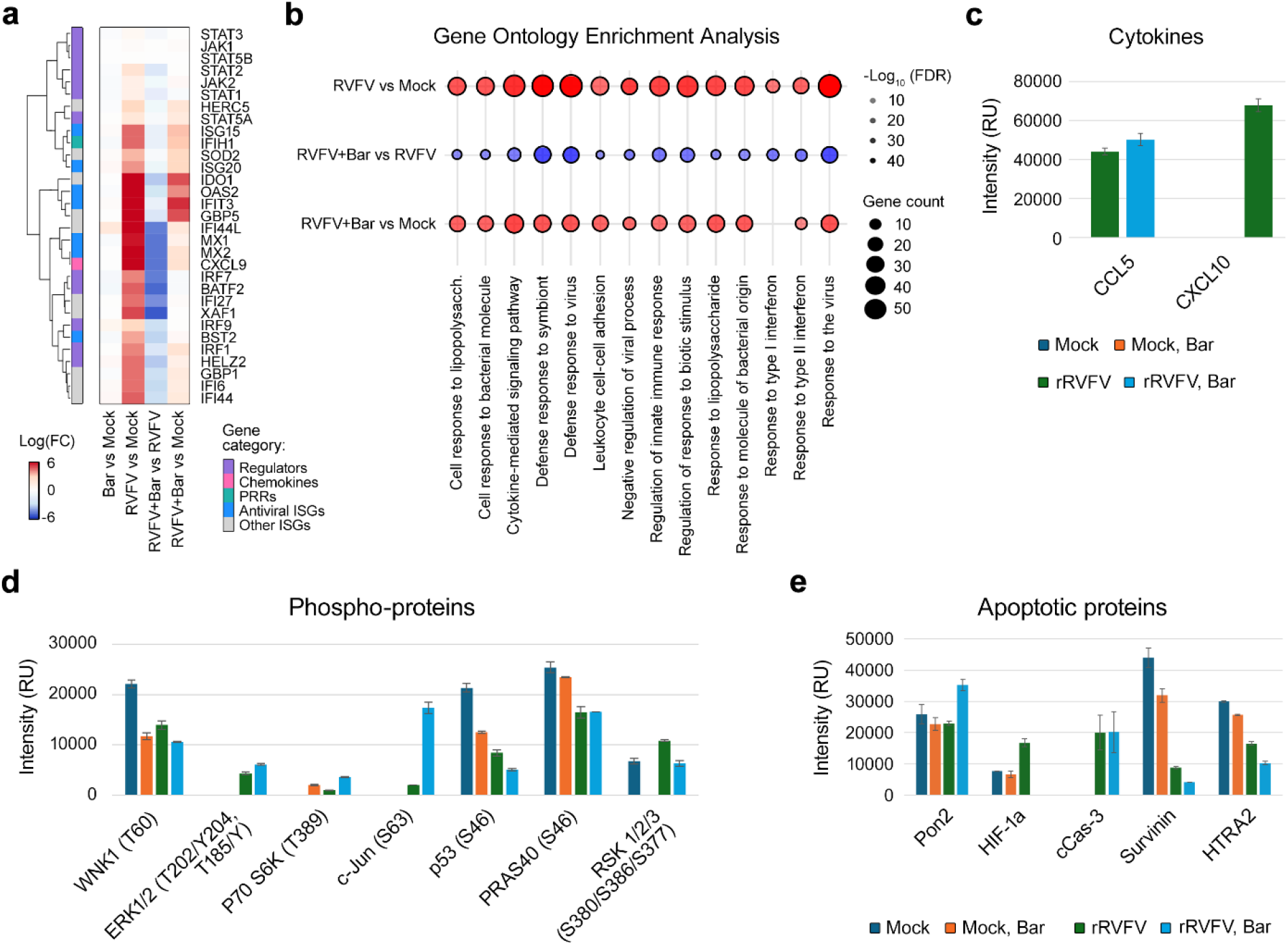
Baricitinib supresses transcription of ISGs, and imbalances signaling and apoptosis in rRVFV-infected A549 cells. (a) Cells were treated with 5 µM baricitinib or vehicle, and infected with the rRVFV or mock. After 8 h, RNA was extracted from cells and sequenced (Mean ± SD, n=3). Differential gene expression analysis was performed. Heatmap showing log_2_ fold changes (logFC) in expression of key interferon response genes across four experimental comparisons. Genes are clustered hierarchically based on expression patterns and annotated by functional categories. Columns represent distinct comparisons: “RVFV vs Mock” shows virus-induced changes; “Bar vs Mock” shows drug effects in uninfected cells; “RVFV+Bar vs RVFV” shows how the drug modulates infection response; and “RVFV+Bar vs Mock” represents the combined effect of virus and drug compared to control. (**b**) Bubble plot representing pathway enrichment analysis for three comparisons (rRVFV vs. Mock, rRVFV/Bar vs. rRVFV, and rRVFV/Bar vs. Mock) is shown. Bubble size indicates the number of genes associated with each pathway (gene count), and bubble color represents the statistical significance of enrichment, expressed as -log10(adjusted p-value). Red bubbles indicate higher pathway activity (upregulation), while blue bubbles suggest lower activity (downregulation or distinct regulation patterns). (**c**) Cells were treated as in (a). After 24 hours, cell culture supernatants were collected, and cytokines were analyzed using the human XL cytokine array; the altered protein levels were plotted (n=2). (**d, e**) Cells were treated as in (a). After 24 hours, cells were lysed; apoptotic and signaling proteins were analyzed using the apoptosis and the phospho-kinase arrays, respectively; the altered protein levels were plotted (n=2).

The cytokine array showed rRVFV-induced production of pro-inflammatory cytokines CCL5 and CXCL10, with baricitinib enhancing CCL5 while completely suppressing CXCL10 (Fig. 4c, Fig. S8a). The phospho-kinase array showed that rRVFV activated MAPK signaling via ERK1/2 and c-Jun phosphorylation while suppressing p53 (S46), reducing apoptosis. Baricitinib amplified ERK1/2 and c-Jun activation but decreased survival-associated kinases such as WNK1 and RSK1/2/3 (Fig. 4d, Fig. S8b). The apoptosis array revealed that rRVFV modulated cell death by increasing HIF-1a and cleaved Caspase-3 while reducing anti-apoptotic Survivin and HTRA2. Baricitinib partially reversed pro-apoptotic changes, suppressing HIF-1a and cleaved Caspase-3, but further reduced Survivin and HTRA2, indicating selective modulation of apoptosis during infection (Fig. 4e, Fig. S8c).

These findings highlight baricitinib’s selective modulation of immune, apoptotic, and signaling pathways, amplifying MAPK activity while dampening IFN responses and survival pathways during RVFV infection.

### 3.5. Baricitinib has pro-rRVFV effect in other cell types and organoids

RVFV infects multiple organs, including the lung, eye and brain, reflecting its ability to target diverse cell types [22,45]. To investigate the pro-viral effects of baricitinib, experiments were conducted using other lung epithelial cells, retinal and brain cells as well aseye and brain organoids.

In lung Calu-3, retinal RPE and brain GBP-1 cells, baricitinib or vehicle was added 15 min prior to infection with rRVFV or mock. Baricitinib enhanced viral replication in a dose-dependent manner, with significant effects observed, without detectable toxicity up to 10 µM (Fig. S2b-d).

Given RVFV’s ability to infect RPE and GBP-1 cells, experiments were extended to retinal and glioblastoma organoids. hROs and hGBOs were treated with 5 μM baricitinib or vehicle, followed by rRVFV or mock infection. After 48 hours, fluorescence imaging showed that baricitinib substantially increased RVFV infection, as demonstrated by heightened Katushka reporter protein expression (Fig. 5a, d; Fig. S9).

**Figure 5.**
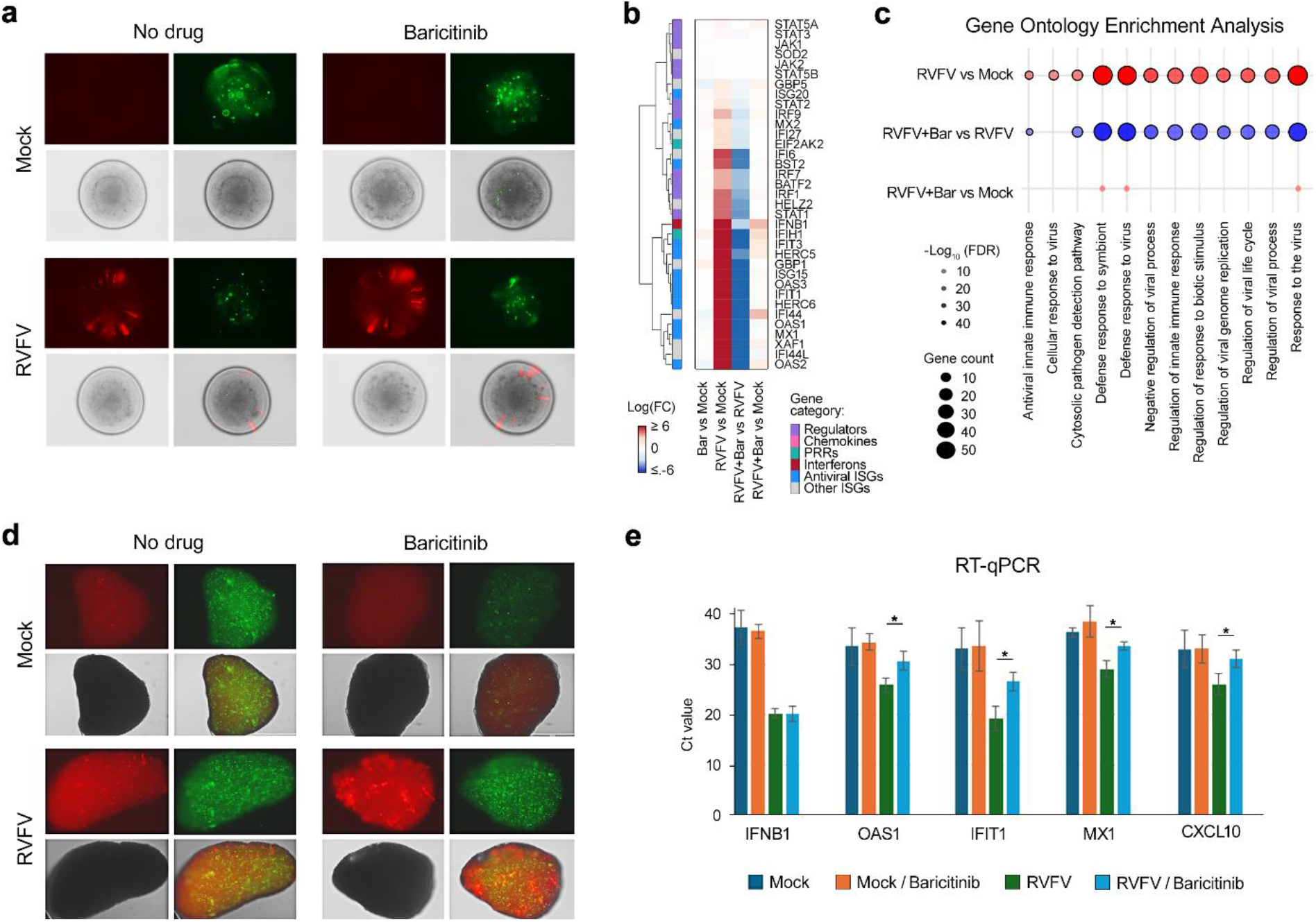
Effect of baricitinib on retinal (hROs) and brain (hGBOs) organoids. (**a**) hROs were treated with 5 µM baricitinib or vehicle, and infected with the rRVFV or mock. After 48 h, microscopic images of organoids were taken. Red channel –far-red fluorescent protein Katushka, green channel – CellToxGreen. Scale bar, 400 mm. (**b**) Organoids were treated as in (a) (Mean ± SD, n=3). After 24 h, RNA was extracted and sequenced. Differential gene expression analysis was performed. Heatmap showing log_2_ fold changes (logFC) in expression of key interferon response genes across four experimental comparisons. Genes are clustered hierarchically based on expression patterns and annotated by functional categories. Columns represent distinct comparisons: “RVFV vs Mock” shows virus-induced changes; “Bar vs Mock” shows drug effects in uninfected cells; “RVFV+Bar vs RVFV” shows how the drug modulates infection response; and “RVFV+Bar vs Mock” represents the combined effect of virus and drug compared to control. inrRVFV/Vehicle, and rRVFV/Baricitinib vs. Mock is shown. The x-axis denotes significantly enriched pathways, while the y-axis represents the conditions compared. Bubble size corresponds to the number of genes involved in each pathway (gene count), and bubble color indicates the direction of regulation (red for upregulated pathways and blue for downregulated pathways). (**d**) hGBOs were treated with 5 µM baricitinib or vehicle, and infected with the rRVFV or mock. After 48 h, microscopic images of organoids were taken. Red channel – mCherry, green channel – CellToxGreen. Scale bar, 400 mm. (**e**) Organoids were treated as in (d) (Mean ± SD, n=3). After 48 h, RNA was extracted and RT-qPCR analyses was performed. Statistical significance between virus-infected and virus-infected drug-treated hGBOs is indicated as * p < 0.05, determined by Wilcoxon rank-sum test with alternative hypothesis ‘greater than’.

Differential gene expression analysis in hROs at 24 h post-infection revealed a robust upregulation of antiviral genes (e.g., *IFIT1, IFIT2, MX1*, and *OAS2*) and pathways related to antiviral innate immunity and viral replication in rRVFV vs. mock comparisons. Baricitinib treatment suppressed these responses, downregulating key interferon-stimulated genes such as *IFIT1, MX1, STAT1*, and *OAS1* in rRVFV/baricitinib vs. rRVFV comparisons, indicating inhibition of IFN signaling (Fig. 5b,c; Fig. S10).

RT-qPCR analysis of RNA isolated from hGBOs at 48 h post-infection revealed a robust upregulation of *IFNB1* and *IFNB1*-stimulated *IFIT1, MX1, CXCL10* and *OAS1*. Baricitinib treatment suppressed the expression of ISGs but not IFNB1 (Fig. 5e; Fig. S10).

These results demonstrate that baricitinib enhances rRVFV infection while suppressing immune responses in lung-, eye- and brain-derived models. This highlights baricitinib’s pro-viral effects and its potential to modulate type I IFN-mediated antiviral pathways across different cell types and organ systems.

### 3.6. Pro-rIAV, rSARS-CoV-2, and AdV effects of baricitinib

To further investigate the pro-viral effects of baricitinib, A549 or A549-ACE2 cells were treated with 5 µM baricitinib or vehicle and infected with rIAV, rSARS-CoV-2, or AdV at an moi of 0.1. In A549 cells infected with rIAV, fluorescence microscopy revealed a marked increase in GFP expression in baricitinib-treated cells compared to vehicle controls, indicating enhanced influenza virus replication (Fig. 6a). Similarly, in A549-ACE2 cells infected with rSARS-CoV-2, baricitinib treatment resulted in increased mCherry fluorescence, suggesting amplified replication of rSARS-CoV-2 (Fig. 6b). In A549 cells infected with AdV, qPCR analysis showed increased secretion of adenoviruses in the media of baricitinib-treated cells (Fig. 6b). This provides further evidence of baricitinib’s ability to promote viral replication by disrupting type I IFN signaling.

**Figure 6.**
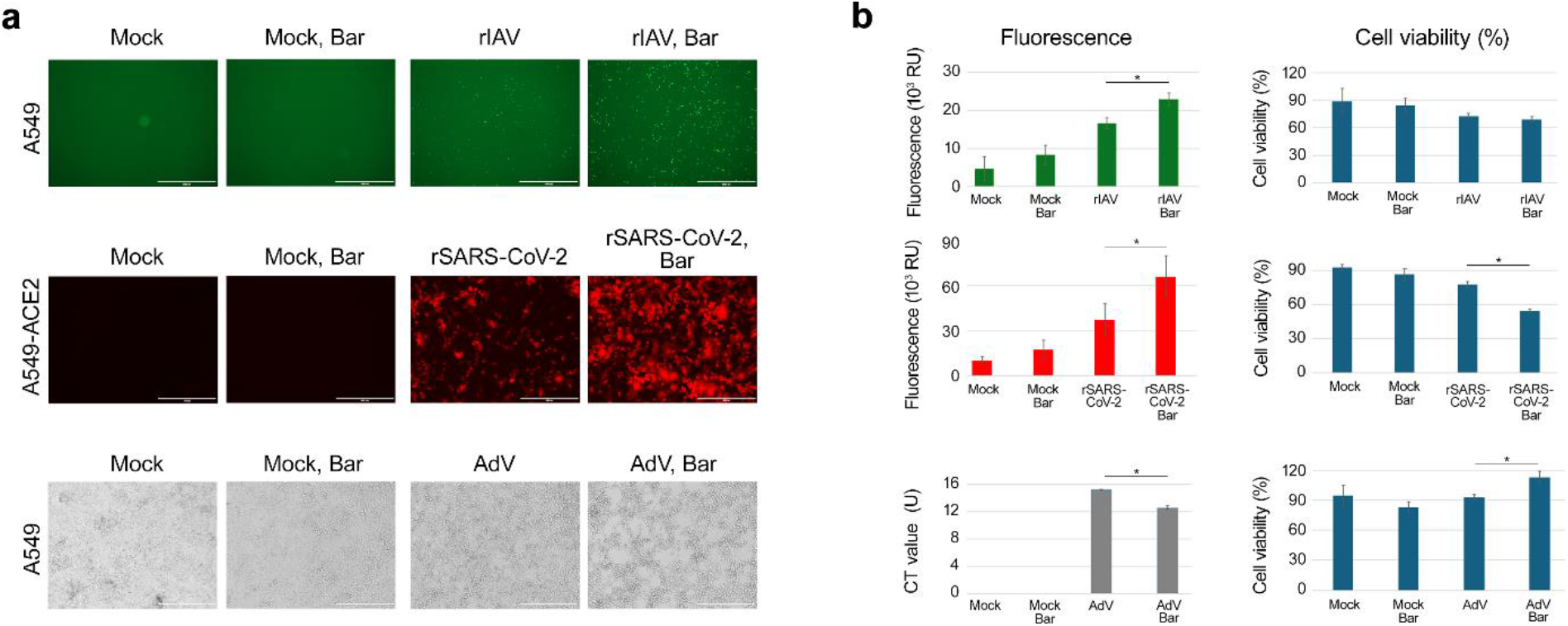
Pro-influenza (rIAV), coronavirus (rSARS-CoV-2), and adenovirus (AdV) effects of baricitinib in A549 or A549-ACE2 cells. (**a**,**b**) Cells were treated with 5 µM baricitinib or vehicle and infected with viruses at an moi of 0.1. After 48 h of rIAV infection, 24 h of rSARS-CoV-2 infection and 72 h of AdV infection images of cells were captured. The red (mCherry) and green (eGFP) fluorescence signals were quantified, normalized and expressed as mean ± SD, (n = 3). Cell viability was assessed using a CTG assay (mean ± SD, n = 3). qPCR analysis of AdV in the media was performed. The CT values were expressed as mean ± SD, n = 3. Cell viability was assessed using a CTG assay (mean ± SD, n = 3). Statistical significance is indicated as * p < 0.05, determined by Wilcoxon rank-sum test with alternative hypothesis ‘greater than’.

## 4. Discussion

Understanding both the therapeutic benefits and potential side effects of medications is crucial. Certain drugs, commonly prescribed for the treatment of other diseases, can inadvertently influence viral infections, potentially altering patient outcomes. For example, anticancer Bcl-xL inhibitors could facilitate killing of animals with virus infections [42,47]. Atorvastatin, candesartan, and hydroxocobalamin could target influenza A virus-host cell interaction and affect the transcription and metabolism of infected cells [46]. JAK inhibitors, while effective in managing inflammation, may increase the risk of viral infections by impairing antiviral defenses and facilitating viral infection [4,6-13,15-21].

In this study, we demonstrated that infection with IFN-sensitive rRVFV induced the synthesis of type I interferons (IFNs) in A549 cells. These IFNs were released extracellularly, bound to IFNR on uninfected cells, and activated the JAK signaling, leading to the production of ISGs. The ISGs provided antiviral protection to the remaining uninfected cells.

However, baricitinib disrupted this protective mechanism by inhibiting JAK signaling, thereby preventing ISG production. This inhibition allowed the virus to spread more effectively. The pro-viral effect of baricitinib was influenced by several factors, including initial viral load, drug concentration, structural characteristics of the inhibitor, cell type, and the addition of other small molecules. Moreover, baricitinib treatment modulated apoptotic and kinase signaling pathways during rRVFV infection in A549 cells. This JAK inhibitor also demonstrated pro-viral effects in retinal and glioblastoma cells and retinal and glioblastoma organoids. Beyond rRVFV, baricitinib enhanced the replication of other viruses, including rIAV, rSARS-CoV-2, and AdV, highlighting a broad pro-viral effect across multiple viruses and cellular models.

Our study has several limitations, as it primarily relies on in vitro models, high drug concentration and includes IFN-sensitive viruses such as rRVFV, rIAV, and rSARS-CoV-2, as well as a single wild-type AdV. These models may not fully capture the complexity of immune responses in vivo. Nevertheless, previous in vitro studies with other viruses, including VSV, RSV, IAV, MeV, MuV, ZIKV, HHV-6A, and ReoV, have also demonstrated that JAK inhibitors can suppress antiviral responses and enhance viral replication [15-21]. Furthermore, in vivo studies have confirmed that JAK inhibition exacerbates infections caused by SARS-CoV-2 and HSV-1. Additionally, clinical evidence suggests that patients with autoimmune conditions who are on JAK inhibitors face an increased risk of hospitalization and death from COVID-19 [12-14], further supporting our findings.

The ability of JAK inhibitors to enhance viral replication presents valuable opportunities in research and biotechnology. Specifically, incorporating JAK inhibitors into cell culture systems can significantly improve diagnostic sensitivity by amplifying low-titer viral samples and boosting the yields of attenuated viruses for vaccine production (EP3801584 and WO2024/003718). Additionally, JAK inhibitors hold promise in antiviral drug discovery as combination agents, enabling viral propagation to identify compounds targeting different stages of viral replication. Furthermore, these inhibitors can enhance the production efficiency of complex oncolytic viruses, achieving higher titers for therapeutic applications. JAK inhibitors can enhance viral vector-based gene delivery by reducing immune responses and improving transduction efficiency. Thus, JAK inhibitors have the potential to revolutionize vaccine production, streamline drug discovery, and improve diagnostic sensitivity for challenging low-titer viral samples.

Understanding the dual nature of JAK inhibitors is critical for optimizing their clinical use. While these drugs provide significant therapeutic benefits in reducing inflammation, they may inadvertently enhance viral propagation by suppressing type I IFN signaling and ISG transcription. Future research should explore the potential for combination therapies that pair JAK inhibitors with antiviral agents to balance immunosuppression and infection risk. Moreover, the structural characteristics identified in this study can guide the design of next-generation JAK inhibitors with reduced pro-viral activity.

Many commonly prescribed drugs could modulate virus-host cell interactions, thus contributing to the morbidity and mortality of patients with virus infections. Further studies are needed to understand the side-effects of the drugs during infectious and other diseases.

In conclusion, JAK inhibitors, such as baricitinib, can suppress type I IFN-mediated antiviral responses, leading to increased viral propagation across various viruses and cell types. This highlights the need for careful consideration when administering these drugs to patients with active viral infections, as they may inadvertently facilitate viral proliferation. Conversely, this property of JAK inhibitors can be harnessed in vaccine development to enhance viral yields in controlled settings, antiviral drug screening in IFN-sensitive cells and organoids as well as in oncolytic virus research to enhance virus-mediated tumor death.

## Institutional review board statement

All experiments with viruses, cells and organoids were performed in BSL2 and BSL3 laboratories in compliance with the guidelines of the national authorities (REK 392418). Standard operational procedures were submitted to the institutional safety committee (Nr. 57317).

## Informed consent statement

Not applicable.

## Authorship contribution statement

Conceptualization and project administration, D.E.K.; methodology, E.R., A.I., J-O.S., D.E.K., M.N.S., H.L., O.V., T.S., H.V., F.W., M.S., M.E., J.F.M., J.E.A., M.B., O.S., W.W., M.K., M.V.; software, A.I.; validation, E.R., M.N.S. and D.E.K.; data curation, A.I., D.E.K., E.R.; writing - original draft preparation, D.E.K, and A.I; results interpretation, D.E.K., M.B.; writing-review and editing, all authors; visualization, A.I.; supervision, M.B., D.E.K., J.E.A., O.V.; funding acquisition, D.E.K., M.B., J.E.A., O.V., and M.E. All authors have read and agreed to the published version of the manuscript.

## Acknowledgements

We thank Andreas Christensen, Annelise Bygland Nikolaisen, Sofie Sagvaag Kristensen, Anette Skjærvik, and Hans-Johnny Schjelderup Nilsen from St. Olavs hospital for clinical samples and RT-qPCR analysis. We thank Knut Jørgen Egelie, Gaute Brede, and Siril Skaret Bakke for their valuable advice on intellectual property rights (IPR).

## Appendix A. Supplementary data

Supplementary Materials: The following are available online: 10 supplementary figures and 2 table.

## References

1. Schwartz, D.M. et al. (2017) JAK inhibition as a therapeutic strategy for immune and inflammatory diseases. Nat Rev Drug Discov 17, 78. 10.1038/nrd.2017.267

2. Tanaka, Y. (2023) A review of Janus kinase inhibitors for the treatment of Covid-19 pneumonia. Inflamm Regen 43, 3. 10.1186/s41232-022-00253-3

3. Jamilloux, Y. et al. (2019) JAK inhibitors for the treatment of autoimmune and inflammatory diseases. Autoimmun Rev 18, 102390. 10.1016/j.autrev.2019.102390

4. Kirito, K. (2023) Recent progress of JAK inhibitors for hematological disorders. Immunol Med 46, 131–142. 10.1080/25785826.2022.2139317

5. Gadina, M. (2024) JAK inhibitors: Is specificity at all relevant? Semin Arthritis Rheum 64S, 152327. 10.1016/j.semarthrit.2023.152327

6. Calvet, X. et al. (2021) Risk of infection associated with Janus Kinase (JAK) inhibitors and biological therapies in inflammatory intestinal disease and rheumatoid arthritis. Prevention strategies. Gastroenterol Hepatol 44, 587–598. 10.1016/j.gastrohep.2021.01.007

7. Boyadzhieva, Z. et al. (2022) Effectiveness and Safety of JAK Inhibitors in Autoinflammatory Diseases: A Systematic Review. Front Med (Lausanne) 9, 930071. 10.3389/fmed.2022.930071

8. Chen, M. et al. (2024) Herpes virus reactivation induced by abrocitinib: A real-world pharmacovigilance analysis of the FDA Adverse Event Reporting System (FAERS) database. Diagn Microbiol Infect Dis 110, 116546. 10.1016/j.diagmicrobio.2024.116546

9. Li, L. et al. (2024) Rapid Activation of VZV After Oral Abrocitinib in a Patient with Atopic Dermatitis. Dermatitis 35, 531–532. 10.1089/derm.2023.0224

10. Pan, C. et al. (2023) Hepatitis B virus reactivation associated with Janus kinase (JAK) inhibitors: a retrospective study of pharmacovigilance databases and review of the literature. Expert Opin Drug Saf 22, 469–476. 10.1080/14740338.2023.2181339

11. Choi, W. et al. (2022) Safety of JAK inhibitor use in patients with rheumatoid arthritis who developed herpes zoster after receiving JAK inhibitors. Clin Rheumatol 41, 1659–1663. 10.1007/s10067-022-06096-0

12. Tsai, J.J. et al. (2023) COVID-19 outcomes in patients with rheumatoid arthritis with biologic or targeted synthetic DMARDs. RMD Open 9. 10.1136/rmdopen-2023-003038

13. Sakai, H. et al. (2024) JAK inhibition during the early phase of SARS-CoV-2 infection worsens kidney injury by suppressing endogenous antiviral activity in mice. Am J Physiol Renal Physiol 326, F931–F941. 10.1152/ajprenal.00011.2024

14. Krzyzowska, M. et al. (2022) Tofacitinib Treatment in Primary Herpes Simplex Encephalitis Interferes With Antiviral Response. J Infect Dis 225, 1545–1553. 10.1093/infdis/jiac040

15. Felt, S.A. et al. (2017) Ruxolitinib and Polycation Combination Treatment Overcomes Multiple Mechanisms of Resistance of Pancreatic Cancer Cells to Oncolytic Vesicular Stomatitis Virus. J Virol 91. 10.1128/JVI.00461-17

16. Gavegnano, C. et al. (2017) Jak Inhibitors Modulate Production of Replication-Competent Zika Virus in Human Hofbauer, Trophoblasts, and Neuroblastoma cells. Pathog Immun 2, 199–218. 10.20411/pai.v2i2.190

17. Islam, S. et al. (2021) Targeting JAK/STAT Signaling Antagonizes Resistance to Oncolytic Reovirus Therapy Driven by Prior Infection with HTLV-1 in Models of T-Cell Lymphoma. Viruses 13. 10.3390/v13071406

18. Schnepf, D. et al. (2021) Selective Janus kinase inhibition preserves interferon-lambda-mediated antiviral responses. Sci Immunol 6. 10.1126/sciimmunol.abd5318

19. Stewart, C.E. et al. (2014) Inhibitors of the interferon response enhance virus replication in vitro. PLoS One 9, e112014. 10.1371/journal.pone.0112014

20. Kim, D. et al. (2020) Targeted therapy guided by single-cell transcriptomic analysis in drug-induced hypersensitivity syndrome: a case report. Nat Med 26, 236–243. 10.1038/s41591-019-0733-7

21. Patel, M.R. et al. (2019) JAK/STAT inhibition with ruxolitinib enhances oncolytic virotherapy in non-small cell lung cancer models. Cancer Gene Ther 26, 411–418. 10.1038/s41417-018-0074-6

22. Ikegami, T. and Makino, S. (2011) The pathogenesis of Rift Valley fever. Viruses 3, 493–519. 10.3390/v3050493

23. Wilson, L.R. and McElroy, A.K. (2024) Rift Valley Fever Virus Encephalitis: Viral and Host Determinants of Pathogenesis. Annu Rev Virol 11, 309–325. 10.1146/annurev-virology-093022-011544

24. Islam, M.K. et al. (2016) High-Throughput Screening Using a Whole-Cell Virus Replication Reporter Gene Assay to Identify Inhibitory Compounds against Rift Valley Fever Virus Infection. J Biomol Screen 21, 354–362. 10.1177/1087057115625184

25. Kittel, C. et al. (2004) Rescue of influenza virus expressing GFP from the NS1 reading frame. Virology 324, 67–73. 10.1016/j.virol.2004.03.035

26. Rihn, S.J. et al. (2021) A plasmid DNA-launched SARS-CoV-2 reverse genetics system and coronavirus toolkit for COVID-19 research. PLoS Biol 19, e3001091. 10.1371/journal.pbio.3001091

27. Chang, C.W. et al. (2022) A Newly Engineered A549 Cell Line Expressing ACE2 and TMPRSS2 Is Highly Permissive to SARS-CoV-2, Including the Delta and Omicron Variants. Viruses 14. 10.3390/v14071369

28. Kuwahara, A. et al. (2015) Generation of a ciliary margin-like stem cell niche from self-organizing human retinal tissue. Nat Commun 6, 6286. 10.1038/ncomms7286

29. Ravlo, E. et al. (2025) Synergistic combination of orally available safe-in-man pleconaril, AG7404, and mindeudesivir inhibits enterovirus infections in human cell and organoid cultures. Cell Mol Life Sci 82, 57. 10.1007/s00018-025-05581-4

30. Jacob, F. et al. (2020) Generation and biobanking of patient-derived glioblastoma organoids and their application in CAR T cell testing. Nat Protoc 15, 4000–4033. 10.1038/s41596-020-0402-9

31. Ianevski, A. et al. (2022) Novel Synergistic Anti-Enteroviral Drug Combinations. Viruses 14. 10.3390/v14091866

32. Ianevski, A. et al. (2024) The combination of pleconaril, rupintrivir, and remdesivir efficiently inhibits enterovirus infections in vitro, delaying the development of drug-resistant virus variants. Antiviral Res 224, 105842. 10.1016/j.antiviral.2024.105842

33. Potdar, S. et al. (2020) Breeze: an integrated quality control and data analysis application for high-throughput drug screening. Bioinformatics 36, 3602–3604. 10.1093/bioinformatics/btaa138

34. Yadav, B. et al. (2014) Quantitative scoring of differential drug sensitivity for individually optimized anticancer therapies. Sci Rep 4, 5193. 10.1038/srep05193

35. Ravlo, E. et al. (2023) A fast, low-cost, robust and high-throughput method for viral nucleic acid isolation based on NAxtra magnetic nanoparticles. Sci Rep 13, 11714. 10.1038/s41598-023-38743-0

36. Wu, T. et al. (2021) clusterProfiler 4.0: A universal enrichment tool for interpreting omics data. Innovation (Camb) 2, 100141. 10.1016/j.xinn.2021.100141

37. Desmyter, J. et al. (1968) Defectiveness of interferon production and of rubella virus interference in a line of African green monkey kidney cells (Vero). J Virol 2, 955–961. 10.1128/JVI.2.10.955-961.1968

38. Diaz, M.O. et al. (1988) Homozygous deletion of the alpha- and beta 1-interferon genes in human leukemia and derived cell lines. Proc Natl Acad Sci U S A 85, 5259–5263. 10.1073/pnas.85.14.5259

39. Andersen, P.I. et al. (2020) Discovery and development of safe-in-man broad-spectrum antiviral agents. Int J Infect Dis 93, 268–276. 10.1016/j.ijid.2020.02.018

40. Ianevski, A. et al. (2022) DrugVirus.info 2.0: an integrative data portal for broad-spectrum antivirals (BSA) and BSA-containing drug combinations (BCCs). Nucleic Acids Res. 10.1093/nar/gkac348

41. Bulanova, D. et al. (2017) Antiviral Properties of Chemical Inhibitors of Cellular Anti-Apoptotic Bcl-2 Proteins. Viruses 9. 10.3390/v9100271

42. Ianevski, A. et al. (2020) Chemical, Physical and Biological Triggers of Evolutionary Conserved Bcl-xL-Mediated Apoptosis. Cancers (Basel) 12. 10.3390/cancers12061694

43. Kakkola, L. et al. (2013) Anticancer compound ABT-263 accelerates apoptosis in virus-infected cells and imbalances cytokine production and lowers survival rates of infected mice. Cell Death Dis 4, e742. 10.1038/cddis.2013.267

44. Fu, Y. et al. (2016) JNJ872 inhibits influenza A virus replication without altering cellular antiviral responses. Antiviral Res 133, 23–31. 10.1016/j.antiviral.2016.07.008

45. Deutman, A.F. and Klomp, H.J. (1981) Rift Valley fever retinitis. Am J Ophthalmol 92, 38–42. 10.1016/s0002-9394(14)75905-7

46. Ianevski, A. et al. (2021) Active Components of Commonly Prescribed Medicines Affect Influenza A Virus-Host Cell Interaction: A Pilot Study. Viruses 13. 10.3390/v13081537

47. Kakkola, L. et al. (2013) Anticancer compound ABT-263 accelerates apoptosis in virus-infected cells and imbalances cytokine production and lowers survival rates of infected mice. Cell Death Dis 4, e742. 10.1038/cddis.2013.267

