## Supplementary material for "JAK inhibitors remove innate immune barriers facilitating viral propagation": 10 Figs and 2 Tables

**Supplementary materials**

Supplementary data (1 table and 10 figures) associated with this article can be found in the online version of the manuscript.

**Table S1:** Primers used for qPCR. Reporter and double-quencher dyes used for probe labeling: 5′ 6-carboxyfluorescein/ZEN/3′ Iowa Black FQ (5′6-FAM/ZEN/3′IBFQ).

| Virus | Primer Sequence (5' to 3') | Acc. Nr. | Target Gene |
|---|---|---|---|
| Adenovirus A, B, D | (+)-TCCGTCAAGAGGCCACTC | X73487 | 5'non-coding region |
| | (+)-CCCGTCAAGAGACTACTC | AY601636 | |
| | (−)-CGGAGCTCAGAGAAATCTC | AF099665 | |
| | (−)-GCGGAGGAGAAAACTCT | | |
| | (−)-AGCAAAGTTTAGAGAAAACTCT | | |
| | (−)-GCTGGCAGAGAAAACTCT | | |
| | [6-FAM]ct+Cg+Ctgg+Ca+Ct+Caa[IBFQ][a)] | | |

**Table S2:** Primers used for RT-qPCR of IFNB1 and ISGs.

| Gene | Forward | Reverse |
|---|---|---|
| IFNB1 | CTGCAACCTTTCGAAGCCTT | AAGCCTCCCATTCAATTGCC |
| MX1 | CCTGTCCCTTCAACCCTCAT | AAGCCGATTCTGACTTCCCA |
| OAS1 | TTGCACCACAGCCTATACCA | ACAGTGAGGAAGGTCATGGG |
| CXCL10 | CTGCCTCTCCCATCACTTCC | AAGCAGGGTCAGAACATCCA |
| IFIT1 | CAAATGATCCACCTGCCTCG | TGGTCACATCTGTCACCTCC |

| Inhibitor | Targets | Therapeutic use | Infection enhancer |
|---|---|---|---|
| Baricitinib*,# | JAK2, JAK1, TYK2, JAK3 | RA, COVID-19 | SARS-CoV-2 |
| Upadacitinib*,# | JAK1, JAK2, JAK1 | RA, PA, AD | HBV, CMV, VZV |
| Tofacitinib*,# | JAK3, JAK2, JAK1, TYK2 | RA, PA, UC | HSV-1 |
| Itacitinib # | JAK1, JAK2 | aGVHD* | |
| Abrocitinib*,# | JAK1, JAK2, TYK2, JAK3 | AD | HSV-1, VZV, |
| Delgocitinib # | JAK1, JAK2, JAK3, TYK2 | AD | |
| Oclacitinib # | JAK1, JAK2, JAK3, TYK2 | RA, IBD | |
| Filgotinib # | JAK1 | Pruritus, AD | |
| Momelotinib* | JAK1, JAK2 | RA, UC, CD | |
| Fedratinib*,# | JAK2, JAK1, FLT3 | Myelofibrosis | HIV |
| Pacritinib*,# | JAK3,JAK2, FLT3 | Myelofibrosis | |
| Ritlecitinib* | JAK3, TEK, ITK, BTK, TXK | Alopecia areata | |
| AT9283 # | JAK1, JAK2, AURKA, AURKB | Leukemia | |
| Ruxolitinib*,# | U-PA, JAK1-3, TYK2 | aGVHD | VSV, RSF, IAV, MeV, MuV, ZIKV, HHV-6A, ReoV, AAV |

*-FDA approved, #-used in this study

**Fig. S1**. JAK inhibitors, their targets, primary clinical indications, and viral infections they may promote. *RA* - rheumatoid arthritis; PA - psoriatic arthritis; AD - atopic dermatitis; UC - ulcerative colitis; aGVHD - Graft-versus-host disease; IBD - inflammatory bowel disease.

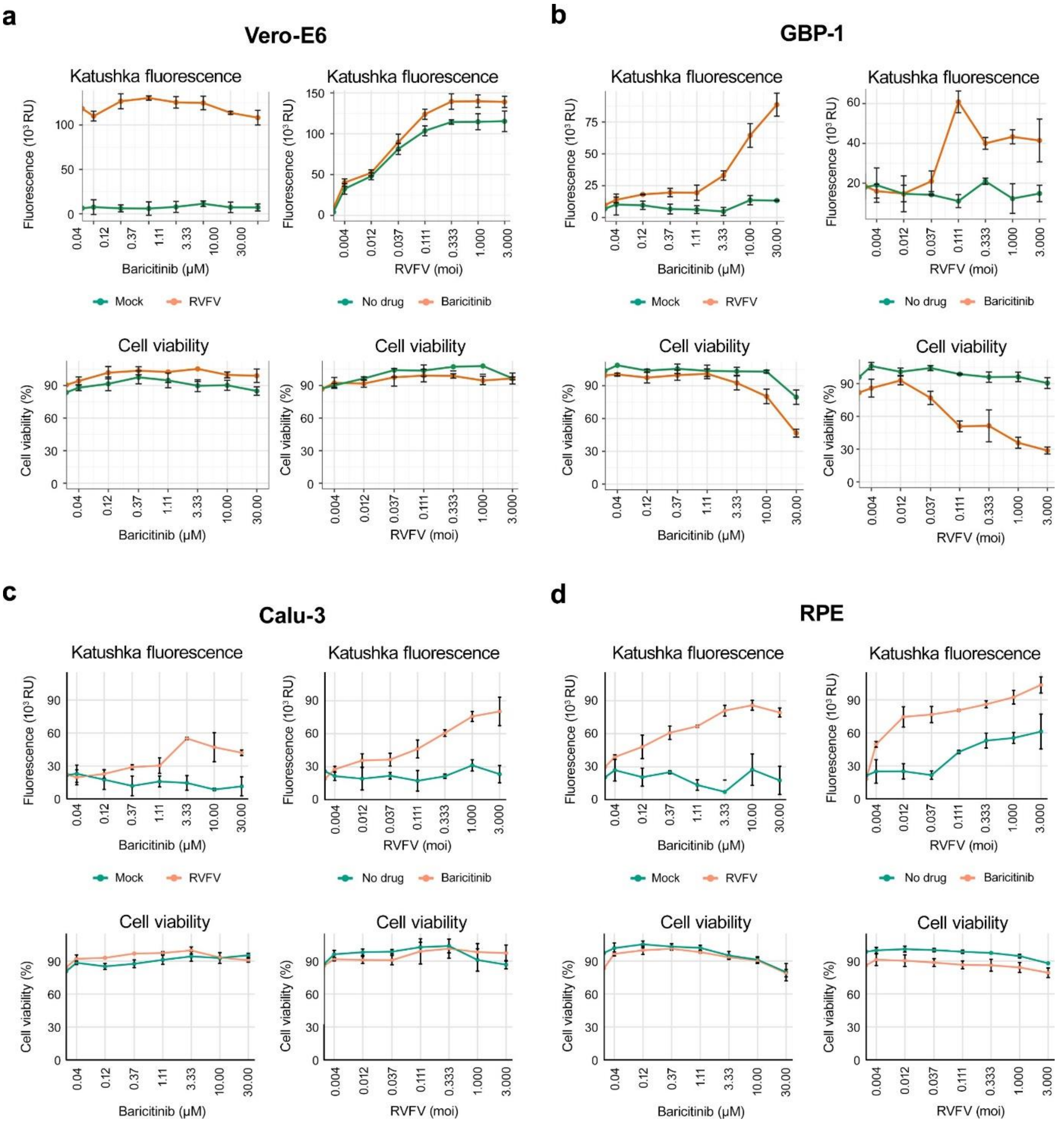


**Fig. S2**. Effect of baricitinib on rRVFV-mediated Katushka expression in Vero-E6, Calu-3, RPE and GBP-1 cells. Cells were treated with vehicle or different concentrations of baricitinib and infected with mock or rRVFV at different moi (moi 0.1 in the experiment with different drug concentration). After 24 (Vero-E6, Calu-3 and RPE) or 48 (GBP-1) hours, Katushka reporter protein expression and cell viability were analyzed (Mean ± SD, n = 3).

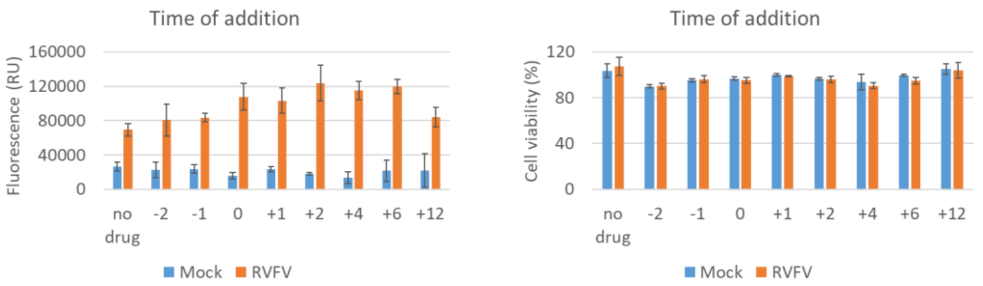


**Fig. S3**. Effect of time of addition of 5 μM baricitinib on rRVFV-mediated expression of Katushka and A549 cell viability (24 hpi).

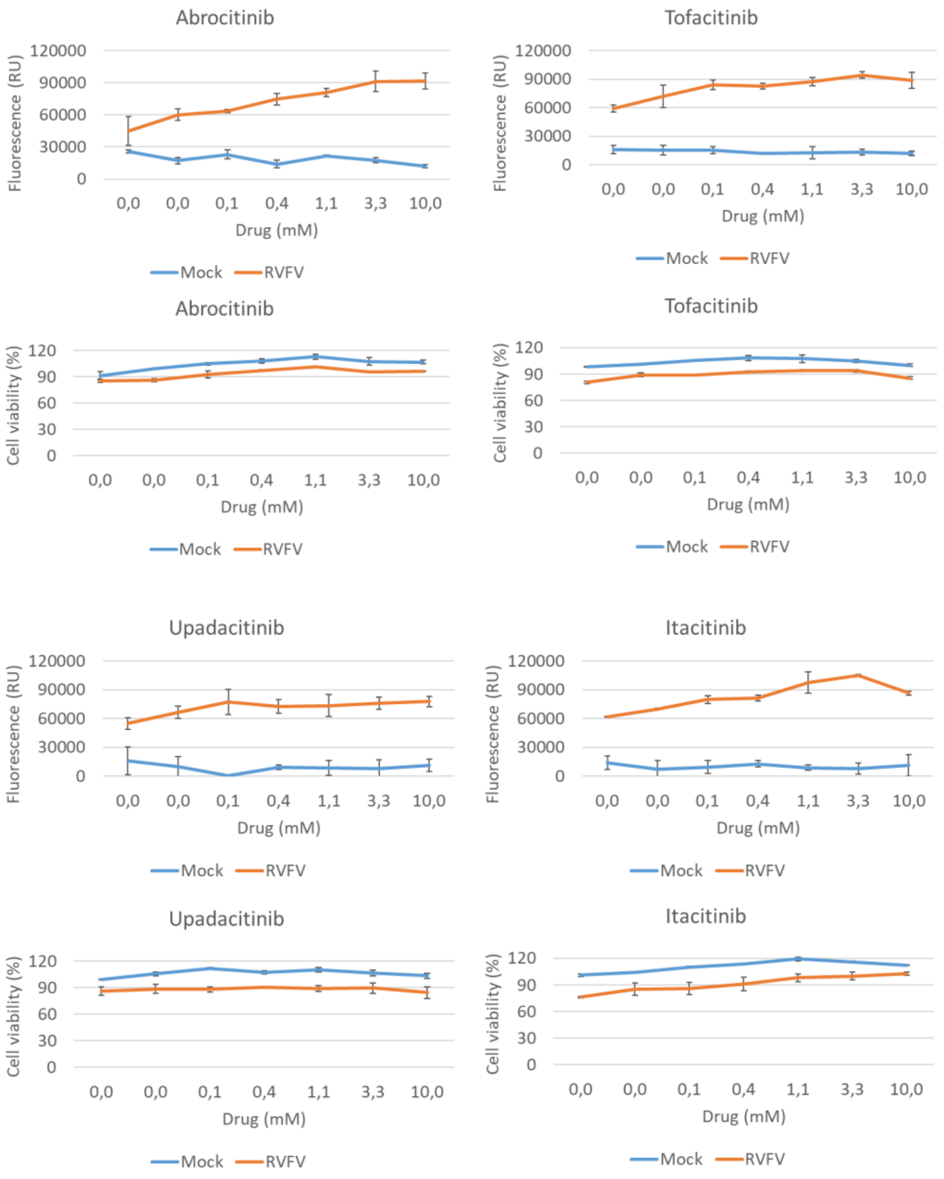


**Fig. S4**. Effect of four Jak inhibitors on rRVFV-mediated Katushka expression and A549 cell viability (24hpi).

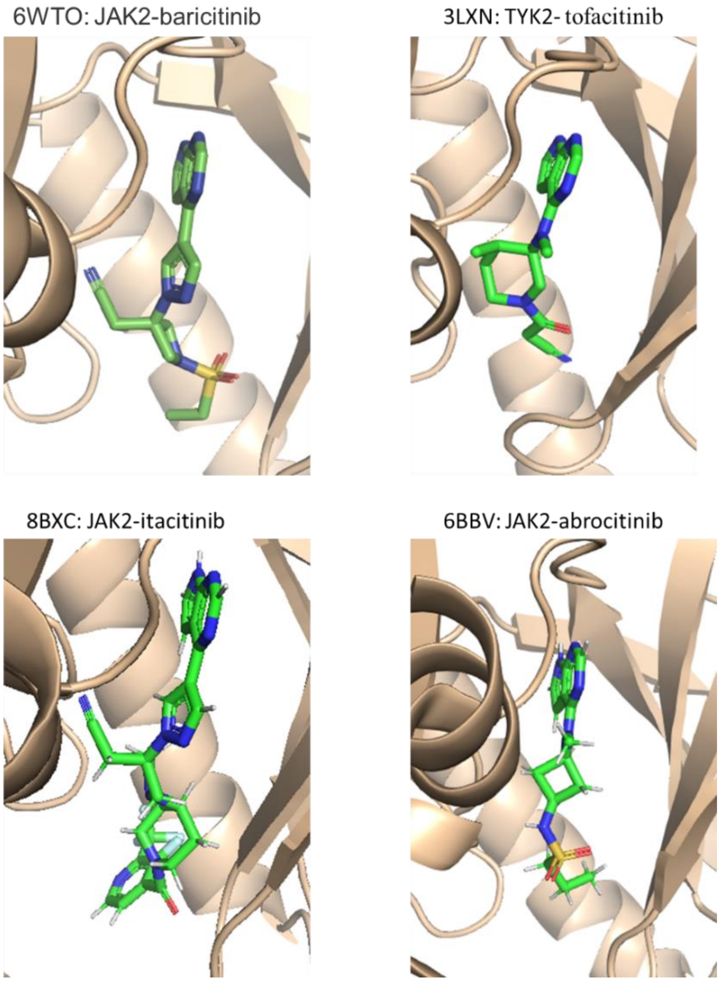


**Fig. S5**. Crystal structures of four inhibitors bound to JAK2 or TYK2, retrieved from the Protein Data Bank (PDB) and visualized using PyMOL. The corresponding PDB accession codes are: 6WTO, 3LXN, 8BXC, and 6BBV.

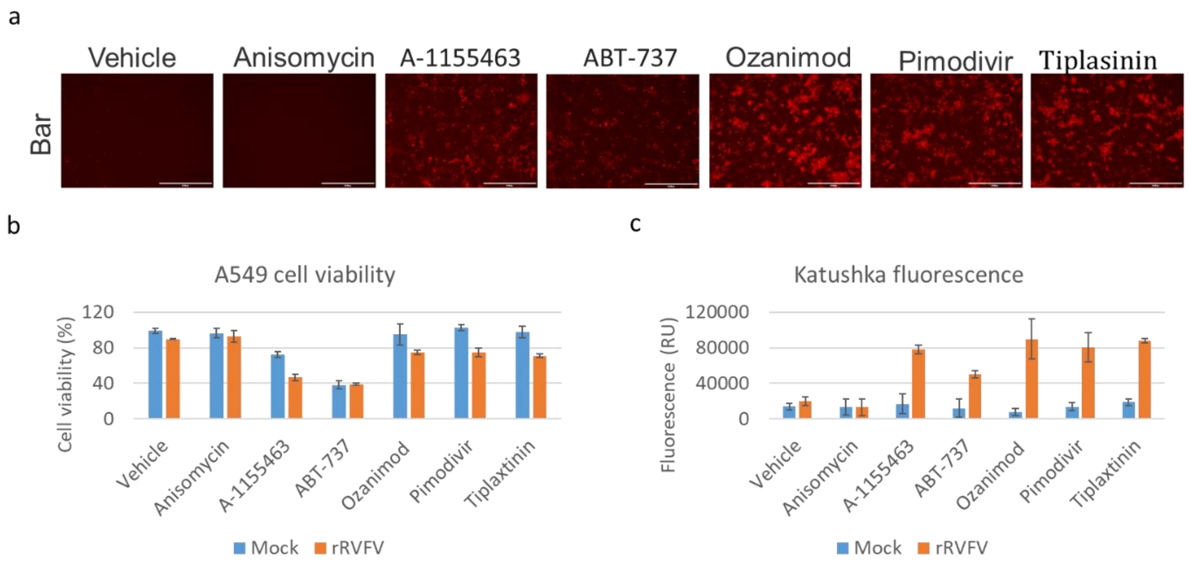


**Fig. S6**. Effect of 10 μM A-1155463, ABT-737, ozanimod, pimodivir, tiplaxtinin, vehicle control and 0,1 μM anisomycin on rRVFV-mediated Katushka expression and viability of baricitinib-treated mock- and rRVFV-infected (moi 0,01) A549 cells (72 hpi).

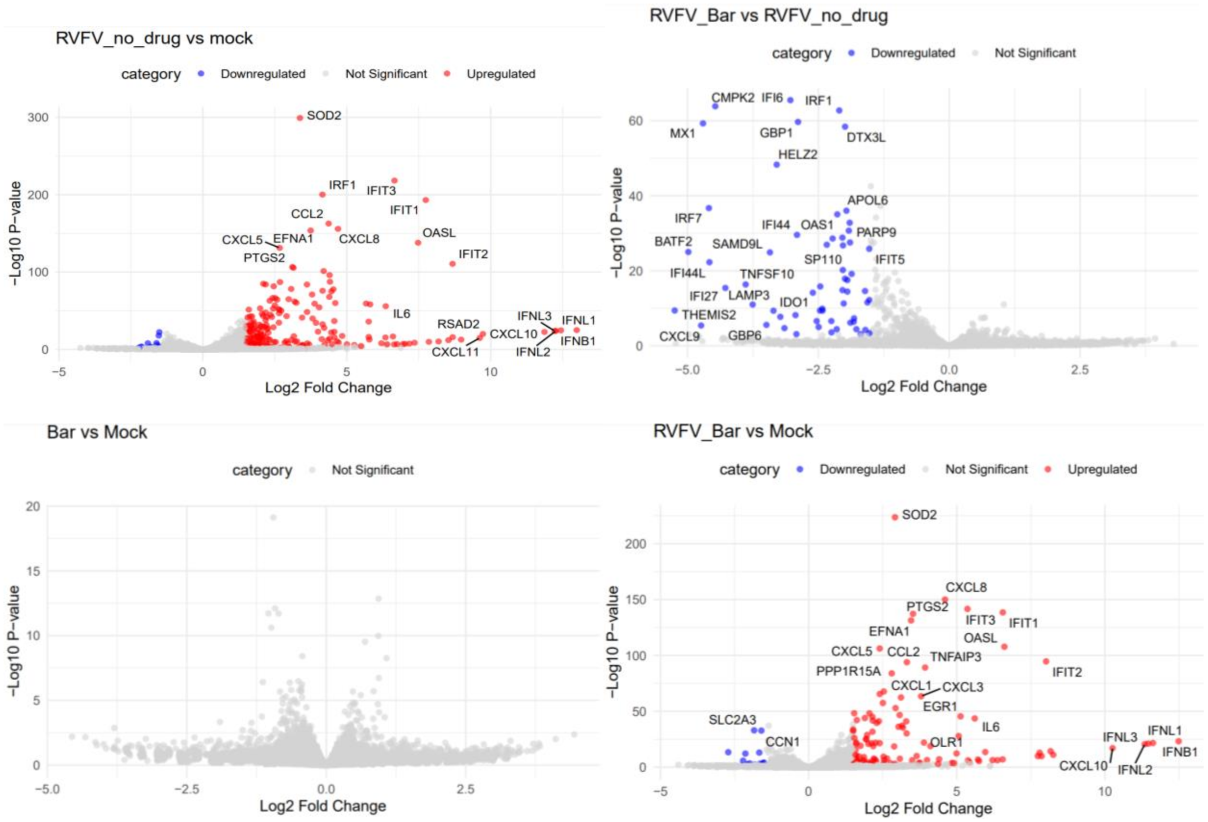


**Fig. S7**. The volcano plots illustrate differential gene expression across four comparisons: RVFV/Vehicle vs. Mock, RVFV/Baricitinib vs. RVFV/Vehicle, Baricitinib vs. Mock, and RVFV/Baricitinib vs. Mock. The x-axis represents the log2 fold change in gene expression, while the y-axis represents the -log10(p-value), indicating statistical significance. Genes are categorized as upregulated (red), downregulated (blue), or not significant (gray).

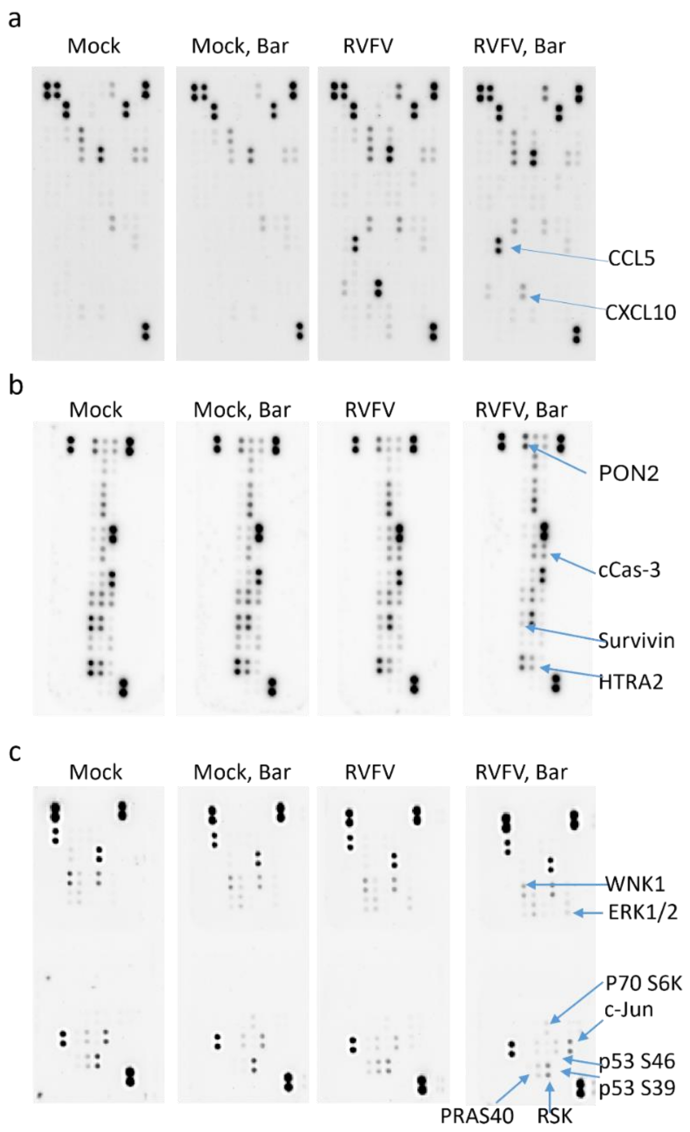


**Fig. S8**. Baricitinib imbalances signaling and apoptosis of rRVFV-infected A549 cells. The cells were treated with 5 µM baricitinib or vehicle, and infected with the rRVFV or mock. After 24 hours, cell culture supernatants were collected, and cytokines were analyzed using the human XL cytokine array; the altered protein levels were plotted (n=2). Remaining cells were lysed; apoptotic and phospoho-proteins were analyzed using the apoptosis and the phospho-kinase arrays, respectively.

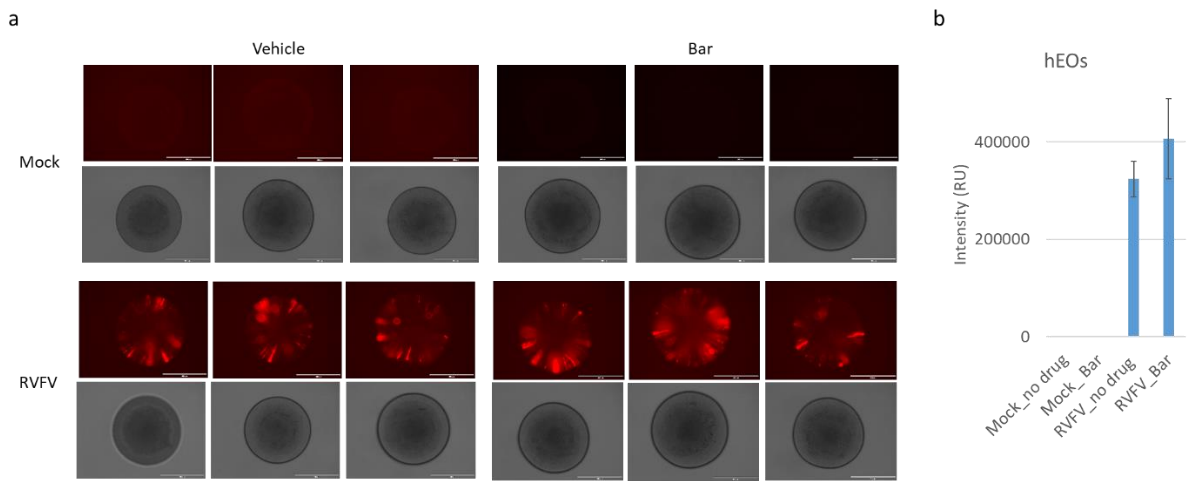


**Fig. S9**. Effect of 5 μM baricitinib on RVFV-mediated Katushka expression in human eye organoids (hEOs; 48 hpi).

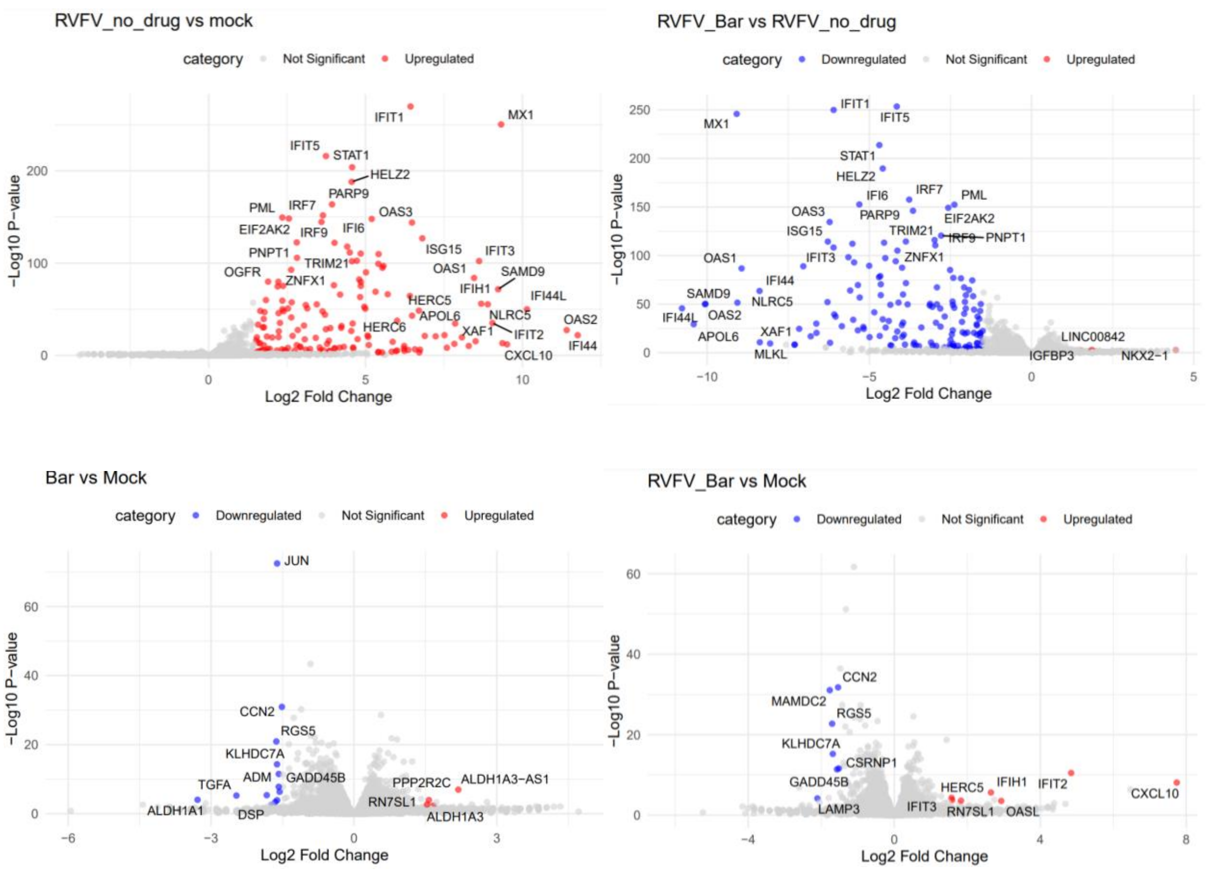


**Fig. S10**. Volcano plots showing differential gene expression in eye organoids under RVFV/Vehicle vs. Mock, RVFV/Baricitinib vs. RVFV/Vehicle, Baricitinib vs. Mock and RVFV/Baricitinib vs. Mock conditions. Upregulated genes are shown in red, and the x-axis represents log2 fold change, while the y-axis represents -log10 (p-value).
